# *tRNAZAP*: Deep learning toolkit for nanopore tRNA signal analysis

**DOI:** 10.64898/2026.09.28.755067

**Authors:** Pooria Daneshvar Kakhaki, Stuart Akeson, Neda Ghohabi Esfahani, Robin L. Abu-Shumays, Miten Jain

**Affiliations:** Department of Electrical & Computer Engineering, Northeastern University, Boston, MA, USA; Department of Bioengineering, Northeastern University, Boston, MA, USA; Biomolecular Engineering Department, University of California, Santa Cruz, CA, USA; Department of Physics, Northeastern University, Boston, MA, USA; Khoury College of Computer Sciences, Northeastern University, Boston, MA, USA

**Keywords:** Sequence-to-sequence Transformer, Ionic current signal segmentation, tRNA classification, Sequence Alignment, Deep learning

## Abstract

tRNA are short, ∼76 nucleotide-long, modification-rich molecules that play an important role in protein translation. Nanopore sequencing is the only platform capable of directly sequencing full-length, native tRNA molecules. This capability is enabled by splinted ligation of tRNA-specific adapters. As a result, the measured ionic current signal reflects a composition of both adapter and tRNA sequence features. Despite these advantages, analysis of nanopore tRNA sequencing data poses distinct computational challenges. Existing ionic current–based signal-to-sequence decoding (basecalling) models are optimized for mRNA and perform suboptimally on short, modification-dense RNA such as tRNA. Furthermore, alignment algorithms for reference-guided analysis of short-length reads are not optimized for nanopore reads with lower accuracy than Next-generation sequencing reads. To overcome these limitations, domain-specific computational approaches are required.

We present ***tRNAZAP***, a nanopore ionic current signal analysis toolkit designed to enable precise reference-guided sequence alignment. tRNAZAP operates directly on raw ionic current signals and decomposes the analysis into two complementary tasks: 1) segmentation of adapter and tRNA regions, and 2) tRNA classification, both implemented using transformer-based models. The segmentation component isolates the tRNA signal portion and extracts the corresponding nucleotide sequence as the alignment query, while the classification component identifies the tRNA type as the alignment reference without requiring sequence-level information. Together, these outputs allow direct optimal pairwise alignment between the tRNA subsequence of the full query and reference sequences. Trained on in vitro transcribed RNA data and evaluated on yeast tRNA, these models achieved ∼98% classification accuracy and intersection-over-union of ∼0.98 for segmentation.

**Graphical abstract:** 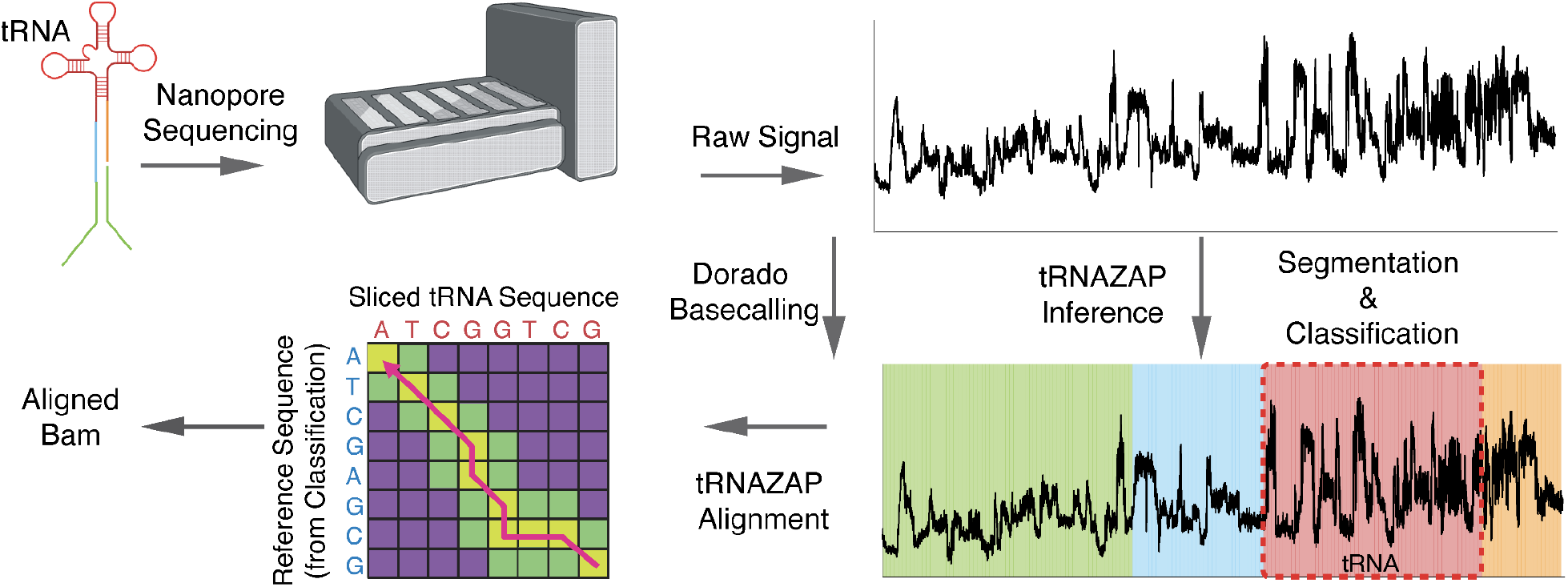

## Introduction

Transfer RNA (tRNA) molecules decode mRNA codons into amino acids during protein translation^1^. Each tRNA contains an anticodon that recognizes specific mRNA codons and delivers the corresponding amino acid to the growing polypeptide chain^1^. tRNA molecules are heavily modified, which affects their translational efficiency and function^2^. Dysregulation of tRNA expression or modification has been implicated in cancer and neurodegenerative disorders, making accurate tRNA characterization essential for understanding translational regulation^3^. However, many of these modifications are lost or misread by sequencing approaches that require reverse transcription or amplification, motivating the use of methods that interrogate native RNA molecules directly^4^.

Nanopore direct RNA sequencing (DRS) enables the analysis of native RNA molecules without the need for reverse transcription^5^. As RNA molecules pass through the nanopore, they disrupt the ionic current in a sequence-dependent manner^6^. The generated ionic current signals, which are recorded as raw electrical signals, reflect both nucleotide sequence and RNA modifications^7,8^. These signals are subsequently decoded into nucleotide sequences by deep learning–based basecalling algorithms. Because tRNA molecules are short in length (typically 76 to 90 nucleotides), they require extension with adapter sequences to enable efficient capture and translocation^9,10^. Splinted ligation methods have been developed to elongate tRNA for nanopore sequencing, thereby producing ionic current signals that contain both tRNA and adapter regions^10,11^.

Recent advances in Nanopore DRS (RNA004 platform and Dorado basecaller) have improved mRNA alignment accuracy to ∼98%^12,13^. However, tRNA basecalling remains challenging due to dense chemical modifications that perturb ionic current signals and confound basecallers trained primarily on longer, less modified RNA sequences^10,11^. This positions raw ionic current as a critical data source for principled tRNA analysis. Here, we present ***tRNAZAP***, a computational toolkit for tRNA-specific nanopore signal analysis that performs classification, segmentation, and alignment directly from raw ionic current signals. By classifying and segmenting signals directly, tRNAZAP enables accurate tRNA identification even when basecalled sequences are unreliable. Our end-to-end deep learning framework uses transformer-based^14^ sequence-to-sequence models to jointly classify tRNA type and segment the signal to isolate the tRNA region from adapter sequences. Segmentation enables the extraction of the query sequence corresponding to the tRNA region. Classification provides candidate reference sequences for the identified tRNA type. Together, these outputs enable direct alignment between query and reference sequences using a cascading alignment strategy featuring an augmented Needleman-Wunsch algorithm^15^ with a fallback Smith-Waterman algorithm^16^. This allowed us to use Nanopore ionic current signal interpretation with sequence-level understanding for robust tRNA alignment and analysis. This signal-level analysis framework can be applied to Nanopore sequencing data for other short RNA species.

Our contributions are summarized below:

- An end-to-end deep learning framework that classifies and segments tRNA raw ionic current signal, providing possible references and query sequences.
- A sequence-alignment algorithm that aligns the query to the candidate reference sequences, measures the distance between them using an augmented Needleman-Wunsch algorithm with a Smith-Waterman fallback.

### Methodology

#### Identification of target sequences

Candidate tRNA sequences for *Saccharomyces cerevisiae* (*S. cerevisiae*) were identified using gtRNA-db 2.0^17^, a product of tRNA-scanSE^18^. Where most tRNA reference sequences are curated to only include a representative subset of sequence diversity^11^, we included the complete set of 79 predicted *S. cerevisiae* tRNA sequences (**Supplementary Table S1**) to demonstrate the resolution of our strategy. This includes 11 sets of tRNA isoacceptors subdivided into 23 isodecoders, a classification of tRNA that often features minimal sequence-level differences. This inclusive set of tRNA encompasses a safe approximation for the possible tRNA classes that could be observed in a biological system, meaning that the model will work for Nanopore tRNA data as long as *S. cerevisiae* tRNA is the input material.

#### *In Vitro* Transcription (IVT) tRNA construction

To train and evaluate *tRNAZAP*, we generated a dataset of *S. cerevisiae* tRNA using *in vitro* transcription (IVT). This required the generation of synthetic DNA templates with a T7 promoter and the reverse complement sequence of our target tRNA. IVT uses biological RNA polymerases to generate target RNA sequences from read template DNA strands. These IVT-derived tRNA molecules allowed us to obtain defined sequence examples without the natural heterogeneity or chemical modifications present in native tRNA. The customizability and cleanliness of the output data make IVT a well-suited strategy for supervised model training.

#### Nanopore library preparation and sequencing

The IVT products were prepared for Oxford Nanopore Technologies (ONT) direct RNA sequencing using a splinted adapter ligation strategy. This required a two-phase molecular preparation: (i) A custom oligonucleotide is added to the 3’-end of the IVT tRNA using T4 DNA ligase. This component is DNA, making the IVT-derived tRNA a hybrid of DNA and RNA. This is in line with our previous methodology of preparing tRNA for sequencing. (ii) After the 3’ DNA oligonucleotide is ligated, we proceed into the RLA ligation step, a standard step in the ONT SQK-RNA004 protocol. The prepared IVT tRNA was sequenced using the ONT SQK-RNA004 chemistry across two PromethION flow cells.

#### Data Preprocessing and Labeling

To obtain reliable training labels, all reads were aligned to the expected *S. cerevisiae* tRNA reference sequences using BWA-MEM^19^ (-x ont2d -w 13 -K 6), which we used to determine the best-matching tRNA class and to assess alignment quality. We selected BWA-MEM over minimap2^20^ for reference alignment because minimap2’s seeding and chaining heuristics are tuned for longer reads and perform inconsistently on tRNA-length sequences (∼76–90 nt)^20^. BWA-MEM has also been the standard aligner in prior tRNA-focused nanopore studies^10,11^. We retained only reads with at least 75% identity to avoid ambiguous or low-quality examples. To create a balanced dataset, we identified the tRNA class with the lowest number of assigned reads (Thr-TGT-1-1, n=44,866) and used the number of reads in that class to determine how many reads to sample from each class: 80% of that amount was assigned to the training set and 10% to the validation set for every class. The rest of the reads were reserved for the test subset.

#### *tRNAZAP* pipeline overview

An overview of tRNAZAP is presented in **Figure 1**. The toolkit consists of two primary components: (1) tRNAZAP Inference and (2) tRNAZAP Alignment. tRNAZAP Inference employs a transformer-based model that jointly performs signal classification and segmentation. Its outputs, together with auxiliary information provided by the basecaller, are subsequently passed to tRNAZAP Alignment, which produces a mapping between the nucleotides of the input tRNA molecule and the corresponding reference sequence. In the following sections, we describe each component in detail and explain how these elements integrate to form the complete tRNAZAP workflow.

**Figure 1.**
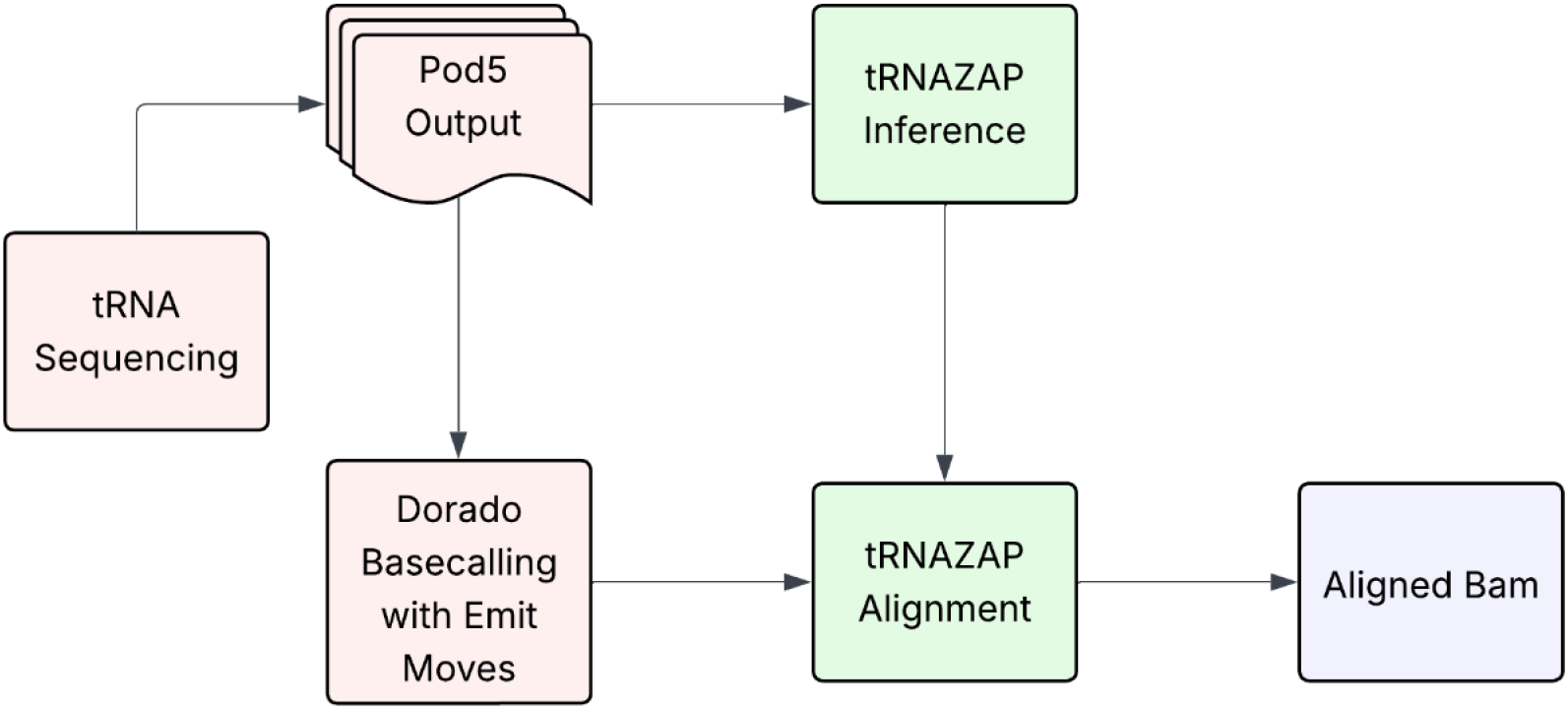
*tRNAZAP* pipeline overview. tRNA sequencing data is processed through Pod5 output and Dorado basecalling with emit-moves. tRNAZAP Inference performs classification and segmentation on raw ionic current signals. tRNAZAP Alignment combines the inference outputs with basecalling information to produce aligned BAM files mapping query tRNA sequences to their corresponding references.

### *tRNAZAP* Inference

We employ a Transformer-based architecture in a multitask learning setup to perform both sequence-level and token-level classification on nanopore ionic current signals. This architectural choice is motivated by the Transformer’s ability to effectively model both local and long-range dependencies in signals, which is required for capturing the hierarchical patterns present in nanopore data. The Transformer model used in this work is encoder-only, as both tasks require processing the input sequence without the need for autoregressive decoding. The architecture includes key components of the standard Transformer, such as stacked layers of multi-head self-attention, position-wise feedforward networks, layer normalization, and residual connections.

The input to the model consists of raw nanopore ionic current signals with an original shape of (*T*, 1), where *T* is the total number of time steps in the signal. Since Transformers operate on sequences of fixed-length tokens rather than continuous streams, we reshape the signal into a sequence of non-overlapping windows, each of size *D*, by splitting it along the time dimension. This results in a sequence of tokens with shape (*T*/*D, D*), where each token represents a window. These windows are then linearly embedded into a higher-dimensional space suitable for the Transformer, producing a sequence of embeddings with shape (*T*/*D, H*), where *H* is the model’s hidden size. A special learnable token, < *CLS* >, is prepended to this sequence to act as a global summary representation for the sequence-level classification task, which results in an input of shape (*T*/*D* + 1, *H*).

To preserve the order of the tokens and enable temporal reasoning, sinusoidal positional encodings are added to the embedded sequence, including the < *CLS* > token. The resulting sequence is then passed to the Transformer encoder, which outputs contextualized representations for both token-level and sequence-level predictions.

The Transformer encoder outputs a sequence of contextualized embeddings with shape (*T*/*D* + 1, *H*), where the first embedding corresponds to the < *CLS* > token and the remaining *T*/*D* embeddings correspond to the tokens derived from breaking the signal into non-overlapping windows.

For **tRNA type classification**, the embedding associated with the < *CLS* > token is passed through a feedforward neural network that serves as the classification head. The final layer of this network contains a number of nodes equal to the total number of tRNA classes, producing one logit per class. These logits are then normalized using a softmax function to generate a score for each tRNA type, representing the model’s estimated degree of class membership.

For **signal segmentation**, the token embeddings associated with windows (*T*/*D* tokens) are each processed independently by a separate classification head. This head assigns each token to one of four categories: ONT adapter, 3′/5′ splint sites, or tRNA. Consequently, the network’s output layer contains only four nodes, each representing one of these regions. Classifying every signal window enables us to subsequently map the predicted labels from window indices back to their respective signal coordinates.

Note that the resulting segmentation is not guaranteed to be strictly contiguous or monotonic across the four classes. Because each window is classified independently, the predicted labels may exhibit local inconsistencies, leading to fragmented or interleaved class assignments (e.g., label sequences such as ONT adapter -> 3′ splint -> tRNA region -> 3′ splint -> tRNA region -> 5′ splint rather than the ideal ONT adapter -> 3′ splint -> tRNA region -> 5′ splint). These artifacts reflect window-level classification noise rather than true biological structure.

To address this, a conditional random field (CRF)^21^ can be applied at the read level to enforce label continuity and ordering constraints. However, performing CRF decoding for every read is computationally expensive and impractical for large-scale analysis. Since our primary focus is the tRNA region, we instead approximate the segment boundaries by defining the start and end positions as the first and last occurrences of that label within each read. This heuristic provides a robust and computationally efficient estimate of the target segment boundaries while avoiding per-read CRF inference. In the rare case where no window receives the tRNA label, segmentation returns a start–end coordinate pair of (-1, -1), and the alignment step falls back to using the entire query sequence rather than a segmented subsequence.

The final outputs of tRNAZAP Inference are therefore twofold: for classification, a score for each tRNA type; and for segmentation, a start–end coordinate pair marking the boundaries of the tRNA region within the raw signal. These outputs are passed to the query sequence extraction step described below.

In our computational experiments, we implemented the transformer encoder with 4 layers, each with 4 attention heads, a 256-dimensional hidden representation, and a 512-dimensional feedforward network. Input signals were divided into non-overlapping windows of 54 timesteps, with a maximum allowance of 1,200 windows. The window size was chosen as a multiple of 6 to remain compatible with the stride of Dorado’s emit-moves output, and the specific combination of window size and maximum window count was selected to fit within our available computational resources. Smaller window sizes are expected to improve segmentation resolution at region boundaries, but proportionally increase the number of tokens per read and the resulting attention matrix size; exploring this trade-off was beyond the computational budget of the present study and is left for future work.

The model was trained independently on the IVT *S. cerevisiae* dataset described in the library preparation section. Ground truth labels for the classification task were derived from the BWA-MEM assignments described in the Data Preprocessing and Labeling section. Ground truth labels for the segmentation task were obtained by mapping BWA-MEM alignment coordinates back to per-timestep positions, yielding a per-timestep region label (ONT adapter, 3′/5′ splint, or tRNA) for each read. These per-timestep labels were then collapsed into per-window labels using a minimum operation over the four class indices (tRNA assigned index 0), such that any window containing at least one tRNA-labeled timestep was assigned the tRNA class. This convention conservatively favors the tRNA label at region boundaries, minimizing the risk of truncating true tRNA signal at the cost of a bounded (≤1 window) inclusion of adjacent non-tRNA signal. Training was performed on four NVIDIA V100 (16 GB) GPUs using data-parallel optimization. Models were trained for 50 epochs with a batch size of 210 signals. Under this setup, each epoch took approximately 20 minutes on average. We optimized the model using AdamW^22^ with a warm-up cosine learning-rate schedule in conjunction with an adaptive gradient–based optimizer tailored for mixed-precision (float16) training.

Two supervised objectives were employed during training. Classification loss *L*_*classification*_ was computed using the cross-entropy between the predicted tRNA class probabilities and the corresponding BWA-MEM-assigned labels. Segmentation loss *L*_*segmentation*_ was defined as the token-wise cross-entropy between predicted window-level labels and the annotated BWA-MEM labels, corresponding to ONT adapter regions, 3′ and 5′ splints, or the tRNA body.

The overall training objective was defined as a weighted sum of the classification and segmentation losses:

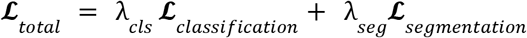

Where *λ*_*cls*_ and *λ*_*seg*_ are non-negative weighting coefficients satisfying *λ*_*cls*_ + *λ*_*seg*_ = 1. These coefficients control the relative contribution of the classification and segmentation objectives during optimization.

Model selection was performed based on validation accuracy for the tRNA classification task, with the best-performing checkpoint retained for downstream evaluation.

An overview of the transformer model is provided in **Figure 2**.

**Figure 2.**
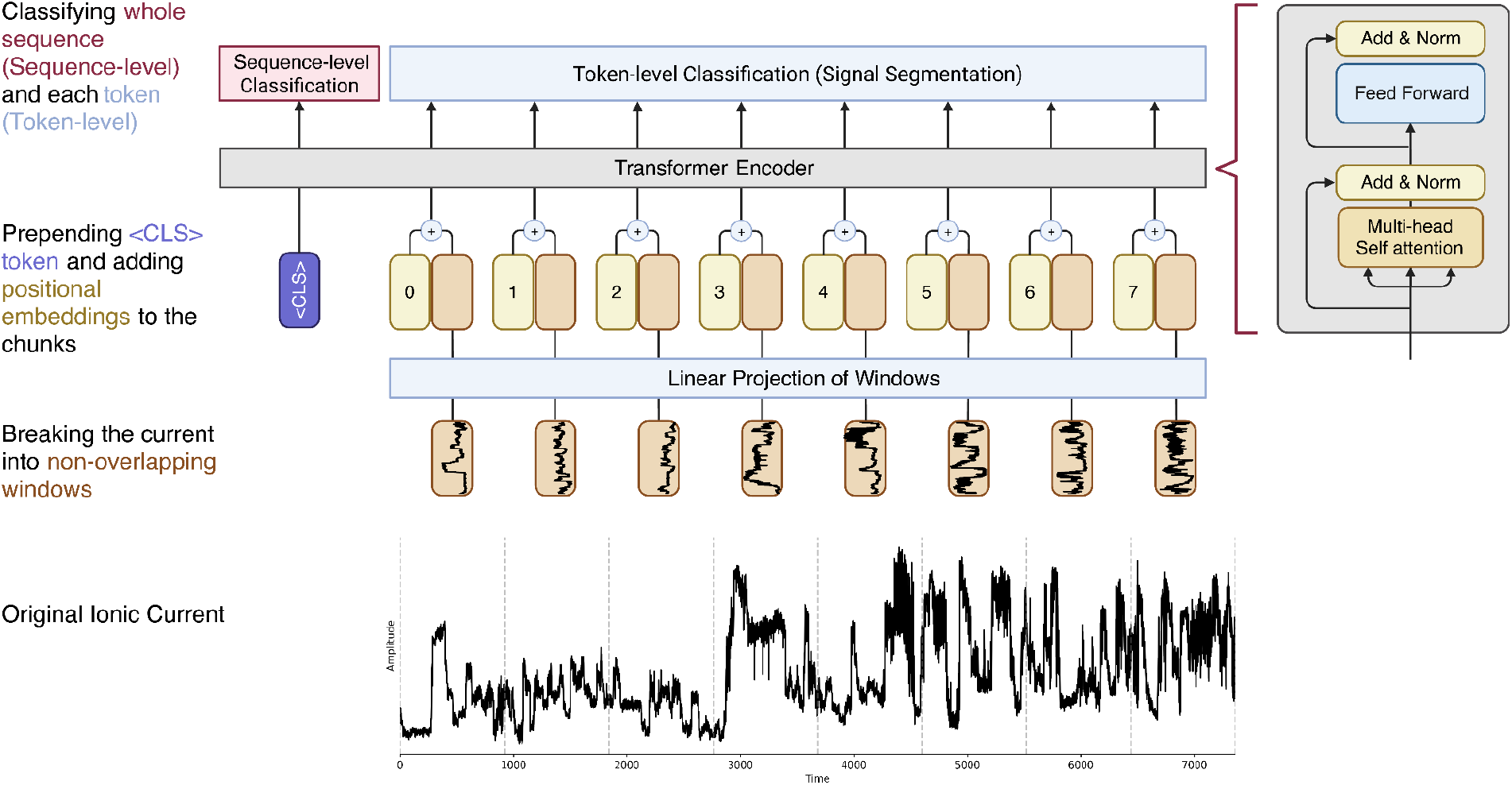
*tRNAZAP* Inference Transformer-based architecture for multitask tRNA signal analysis. Raw ionic current signals are divided into non-overlapping chunks and linearly projected into token embeddings. A < *CLS* > token is prepended, and positional embeddings are added before processing through the transformer encoder. The encoder consists of stacked layers with multi-head self-attention, feed-forward networks, and add & norm operations. The < *CLS* > token output is used for sequence-level classification (tRNA type identification), while individual token outputs are used for token-level classification (signal segmentation into ONT adapter, 3’/5’ splint, and tRNA regions).

**Figure 3.**
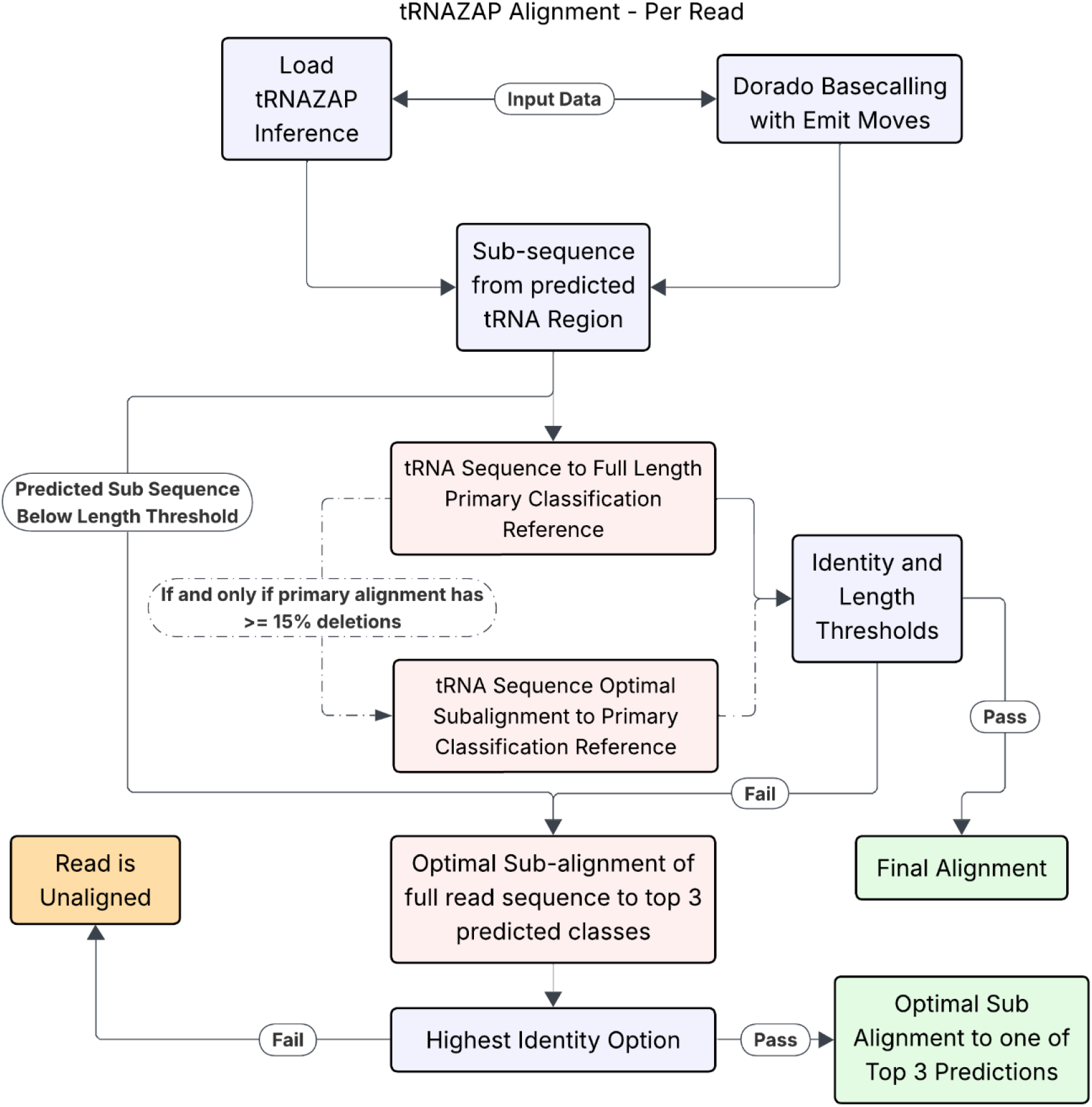
*tRNAZAP* Alignment Strategy. A schematic of the *tRNAZAP* alignment strategy at a per-read level. Pink boxes indicate alignment steps, green boxes are successful alignments, orange boxes are failed alignments, and blue boxes indicate intermediate data preparation and decision steps.

### Query sequence extraction

The classification module of tRNAZAP Inference produces per-tRNA type classification scores that identify the most probable reference sequence for the input signal. These scores provide a ranked set of candidate references, enabling downstream analyses to focus on the most likely tRNA isoacceptor or isodecoder sequences.

The segmentation module complements this by outlining the precise start and end coordinates of the signal belonging to the tRNA molecule. In our workflow, we make use of the Dorado basecaller’s emit-moves functionality (which maps signal positions to basecalled nucleotides^23^). Dorado basecalls the entire raw signal, including adapter and splint regions, and therefore does not by itself yield tRNA-specific sequence information. However, it provides emit-moves, which are a mapping between signal ranges and nucleotide positions. Because tRNAZAP precisely determines which portion of the signal corresponds to the tRNA region, the combination of emit-moves with tRNAZAP’s segmentation enables accurate extraction of the nucleotide bases that belong specifically to the tRNA molecule.

Having identified both the query sequence (extracted from the tRNA-specific segment of the signal) and the most likely reference sequence, the workflow proceeds with alignment. tRNAZAP alignments integrate these inputs to produce a high-resolution mapping between the observed tRNA sequence and the predicted reference, enabling downstream characterization of sequence variation and structural features within the tRNA molecule.

### *tRNAZAP* Alignment

tRNAZAP alignment performs a pairwise alignment between the predicted tRNA portion of the read and the predicted tRNA class reference, both from the inference step. We use a cascading alignment strategy to the top predicted reference that (i) attempts to align the tRNA sequence to the full predicted reference sequence using a modified Needleman-Wunsch algorithm, and (ii) if more than 15% of the alignment is a deletion, falls back to a Smith-Waterman algorithm, calculating the optimal subsequence alignment. (iii) Finally, if neither previous strategy works, we default to a Smith-Waterman alignment of the entire tRNA sequence to each of the top three reference options.

For the Needleman-Wunsch algorithm, we are attempting to align the predicted sequence to the entire associated reference. To this end, we augmented the Needleman-Wunsch algorithm to allow for a truncated alignment of the read to the full reference. To accomplish this, we initialize the dynamic programming matrix with 0 scores in the first column. The dynamic programming matrix is populated with an affine gap penalty scoring scheme. The scores for gap open, gap extend, and mismatch are adjustable parameters. Once the dynamic programming matrix is populated, we select the cell in the final column that has the lowest score to utilize query bases. This relationship can be formalized with the following equation.

If this strategy produces an optimal alignment with a deletion rate of 15% or greater, we consider a shift from a Wagner-Fischer pseudo-global alignment strategy^24^ to a Smith-Waterman optimal sub-alignment strategy. The Smith-Waterman optimal subalignment is performed with independent and adjustable affine gap cost parameters. The decision to switch to Smith-Waterman is made independently for the primary and secondary classification reference. This means it’s possible for a primary Wagner-Fischer alignment to be compared to a secondary Smith-Waterman alignment and vice versa. All of these alignments are performed with the slice of read sequence predicted to correspond to the tRNA portion of the read. Finally, we trim the leading characters of the alignment until a window of 8 nucleotides achieves an alignment accuracy of 75%. This helps to account for nanopore sequencing-specific quality uncertainty towards the 5′ end of individual molecules. By default, tRNAzap penalizes gaps differently for its two alignment modalities. For the Needleman-Wunsch style global alignment gap opening and extensions are scored as 1.0 and 0.7, respectively. For the Smith-Waterman local alignment matches score 3, mismatches -3, gap opening -6, and gap extension -3.

If a suitable alignment cannot be found using these strategies, a final optimal subalignment is performed using the entire read sequence against the top three predicted reference options. This Smith-Waterman based step uses the same affine gap penalties as the Smith-Waterman of the previous step. Across all of the alignment strategies, the condition for a read passing or failing is dependent on meeting a minimum number of matching bases in the alignment and achieving at least a minimum alignment identity. These parameters are adjustable to accommodate a variety of sequencing and alignment conditions. Ultimately, tRNAZAP’s alignment module outputs a standard format BAM file, along with all typical tags associated with Nanopore DRS. This means that tRNAZAP can be directly plugged into a downstream bioinformatic pipeline for processing without any additional data modulation steps.

## Results

### Dataset summary

The sequencing experiment yielded a total of 18,858,466 reads across the 79 classes. Among these, 9,386,667 reads were retained to construct the full dataset after applying the filtering criteria (see Data Preprocessing and Labeling section). **Supplementary Table S2** documents the number of reads per subset for each tRNA class.

### *tRNAZAP* Inference classifies and segments the tRNA raw ionic current signals

An example output of tRNAZAP inference model is shown in **Figure 4**, demonstrating the classification and segmentation components of the tRNAZAP Inference. The segmentation component identifies boundaries between the ONT adapter, 3’ splint, tRNA, and 5’ splint regions from raw ionic current (**Figure 4A**). The classification component then assigns scores to each one of 79 *S. cerevisiae* tRNA classes. (**Figure 4B**).

**Figure 4.**
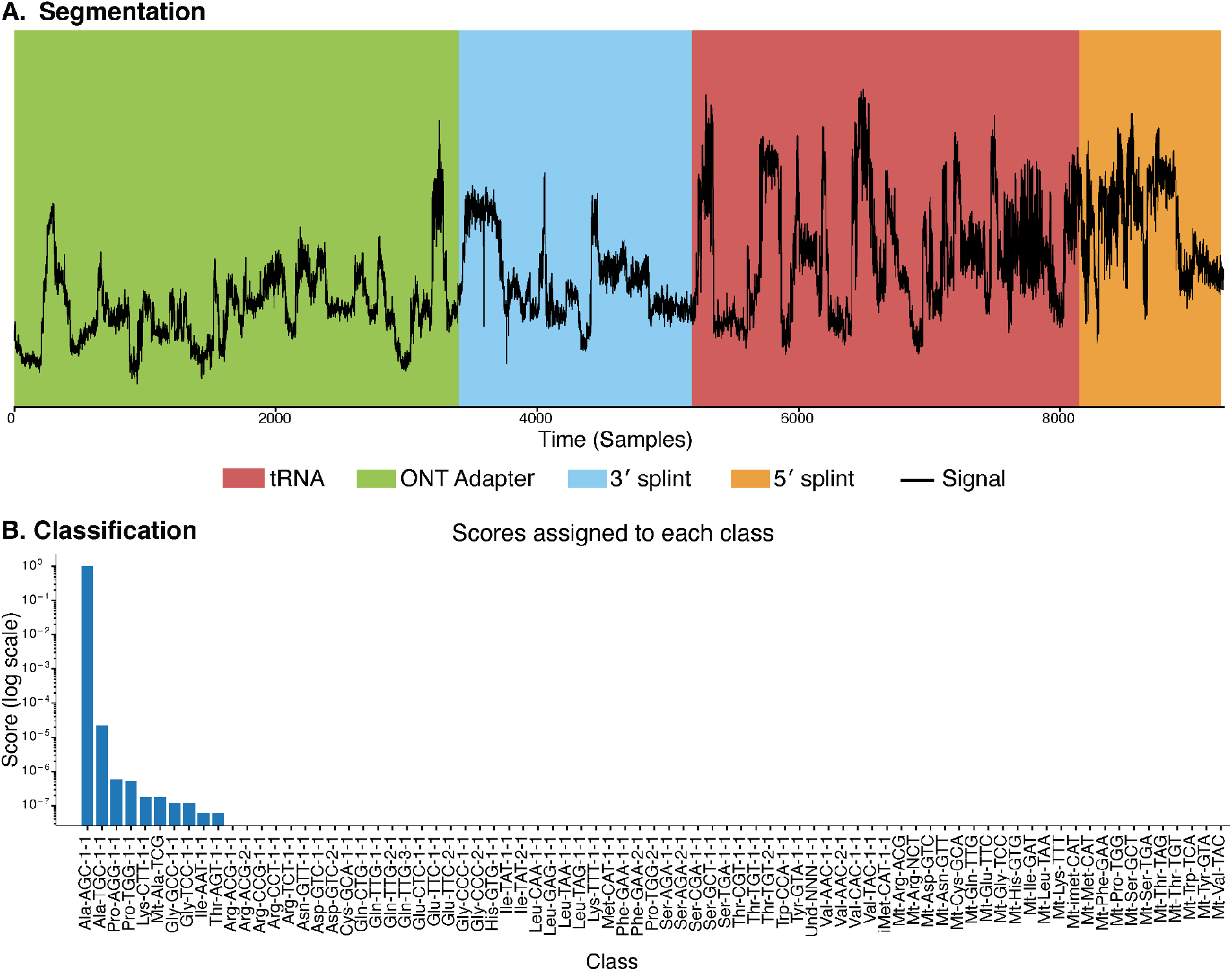
*tRNAZAP* inference example. **A.** Segmentation of ionic current signal into ONT adapter (green), 3’ splint (blue), tRNA (red), and 5’ splint (orange) regions. **B**. Classification scores for a single read across 79 *S. cerevisiae* tRNA classes, with the top true predicted class (AlaAGC-1-1) with highest score.

The model was applied to all reads in the test dataset to evaluate its performance. The classification and segmentation components were assessed independently, with classification performance quantified by accuracy across the whole dataset, precision, recall, and F1 score within each class. Segmentation performance was quantified using the intersection-over-union (IoU) metric, computed between the predicted and BWA-MEM tRNA region ranges.

In the classification task, the model achieved a micro-average-accuracy (overall accuracy across all samples) of 99.05% and a macro-averaged-accuracy (unweighted average accuracy across classes) of 98.83%. Per-class performance was further examined using the confusion matrix, which shows >=89.69% diagonal values indicative of robust class discrimination (**Figure 5.A, Supplementary Figure S1)**. To further assess the classification performance, we examined how the model assigned scores to each read. Specifically, we analyzed the distribution of scores assigned to the ground-truth class for each category. Across all classes, 96.83% of predictions fall within the 0.975–1.0 confidence range. (**Figure 5.C)**. Per-class precision, recall and F1 scores are provided in **Supplementary Table S3**.

**Figure 5.**
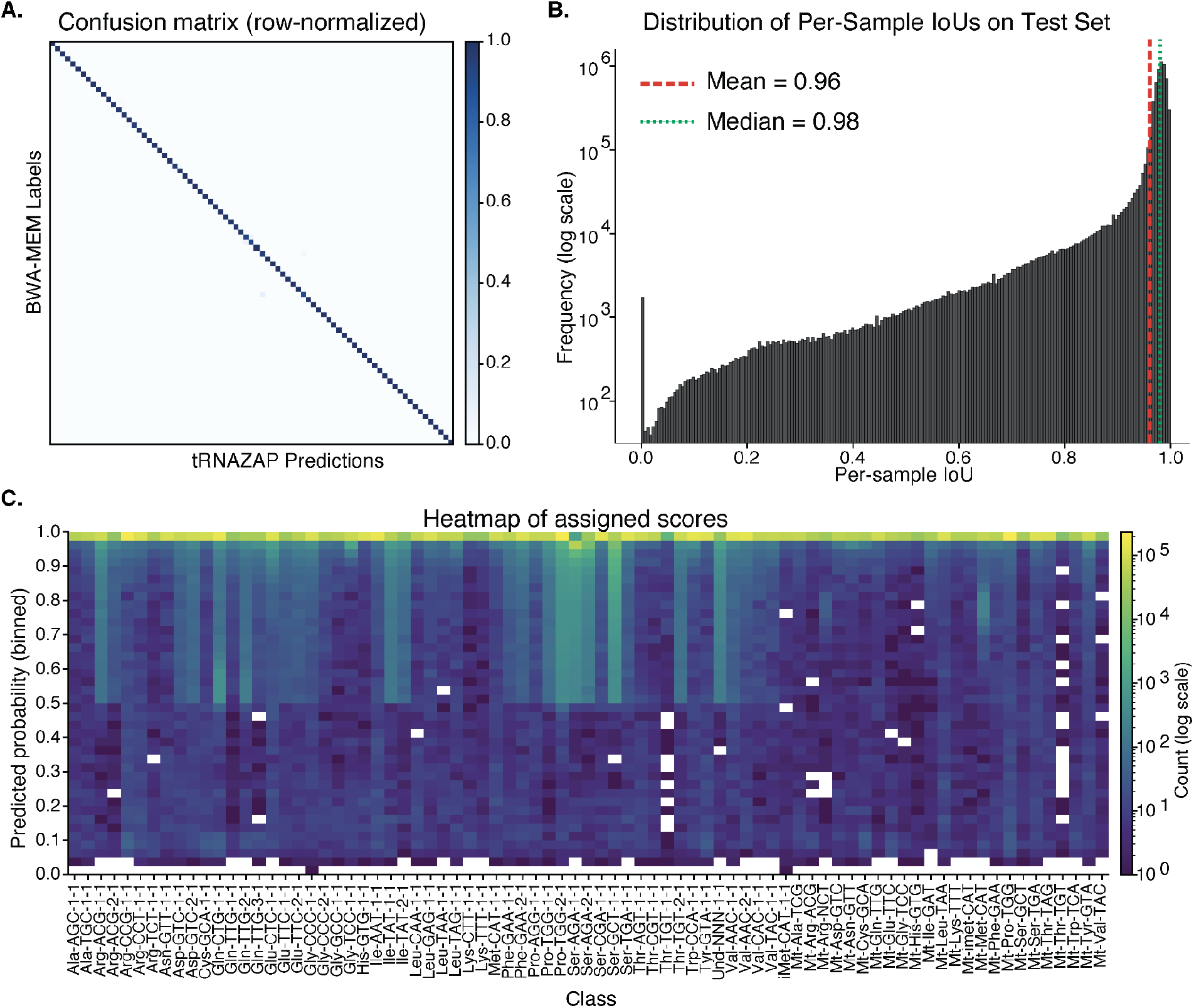
Classification and segmentation performance of *tRNAZAP* on the test set. **A.** Row-normalized confusion matrix showing classification accuracy across 79 *S. cerevisiae* tRNA classes. **B**. Distribution of per-sample intersection-over-union (IoU) scores for segmentation, with mean and median values indicated. **C**. Heatmap of classification scores assigned to each tRNA class across different predicted probability bins, showing the concentration of predictions at high confidence levels.

In the segmentation task, the model achieved an average IoU of 0.96. The distribution of per-sample IoU values is skewed toward the upper range, with a median of 0.98 **(Figure 5.B)**.

### Transformer attention concentrates on the tRNA region

To better understand and interpret the behavior of our transformer model, we analyzed how the < *CLS* > token attends to different regions of the input signal. Because the ONT adapter and the 3′/5′ splint regions are not molecule-specific and are identical across reads, they provide no information relevant to distinguishing tRNA species. In contrast, the tRNA region contains the specific features that determine the read’s class. Therefore, we expect the < *CLS* > token to focus its attention primarily on this region.

Figure 6 demonstrates that the attention profiles of the individual heads, as well as their mean, concentrate predominantly on the tRNA region. This pattern supports our assumption and indicates that the model is relying on biologically relevant ionic current signal features when performing classification.

**Figure 6.**
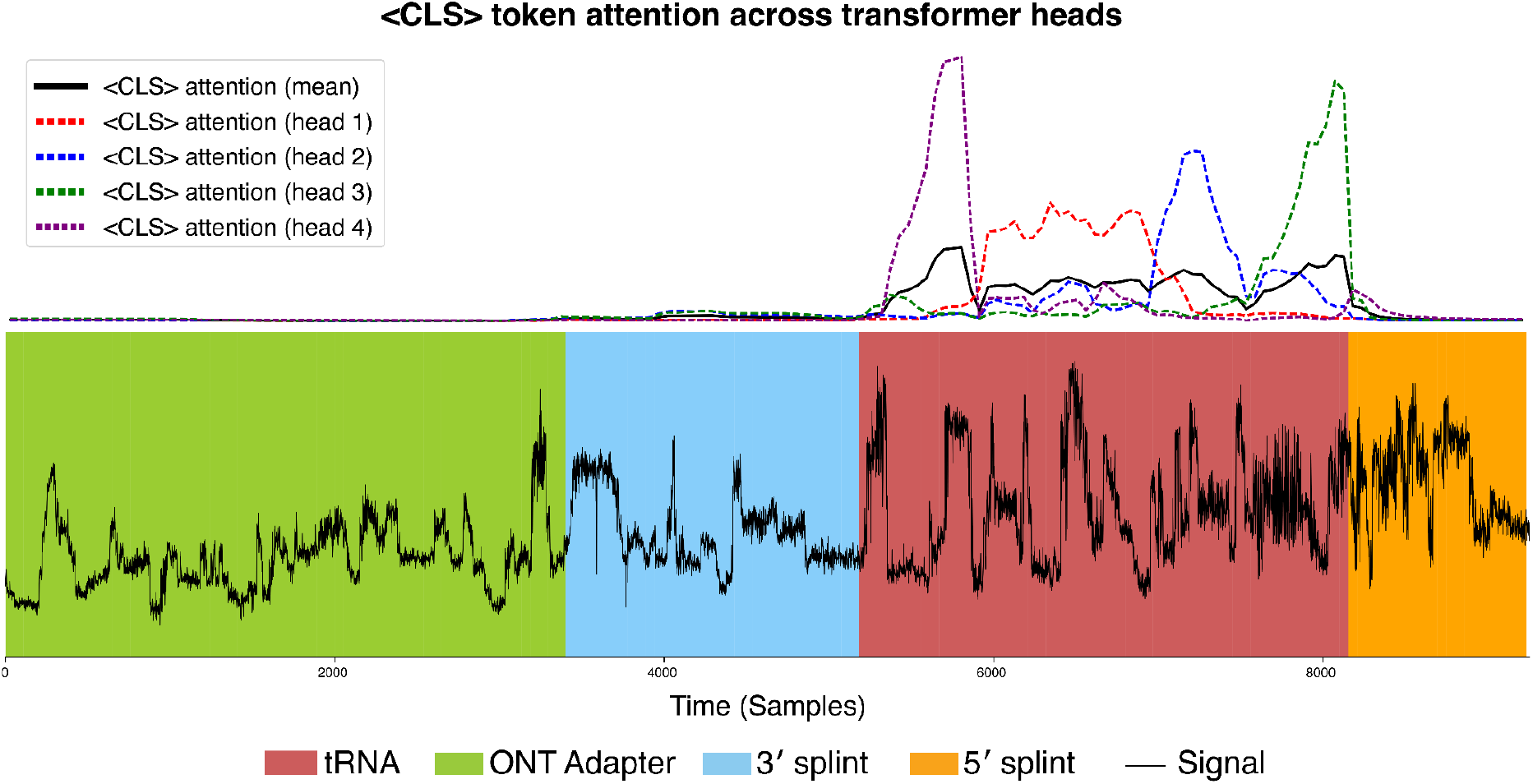
Attention patterns of the < *CLS* > token across transformer heads. Attention weights from each of the four attention heads (colored dashed lines) and their mean (black solid line) are plotted above the segmented ionic current signal. The < *CLS* > token concentrates attention primarily on the tRNA region (red), with minimal attention to the ONT adapter (green), 3’ splint (blue), and 5’ splint (orange) regions.

### *tRNAZAP* Alignment

Contemporary alignment tools for genomic data typically rely on comparing the alignment scores across all available reference sequences. The ability to differentiate between sequences depends on the accuracy of the query sequence and the amount of separation between reference sequences. In the case of biological tRNA both of these factors work against successful alignment of nanopore-derived tRNA read sequences. For all three kingdoms of life, there are examples of tRNAs with a single nucleotide difference across the full reference body. This is further complicated by the dense population of chemical modifications in tRNA that can lower sequence prediction accuracy.

tRNAZAP stands apart from these strategies by predicting the reference class based solely on ionic current, a richer source of information in comparison to sequence-level information. We developed a pairwise alignment strategy to identify the globally optimal alignment of the query sequence to the reference sequence associated with the predicted class. We augmented the Needleman-Wunsch pairwise alignment strategy to allow for affine gap penalties and to minimize the edit distance normalized by the number of aligned bases. Because a prediction of class will be produced for every read, the alignment step had to filter out sub-par alignments, including non-tRNA reads. To this end, we enforced a cutoff of a minimum of 25 matching bases as well as a minimum alignment identity of 75% across the entire aligned region (**Figure 7.A**). When we compared the alignment identity of BWA-MEM aligned reads with tRNAZAP alignment identity, we observed a strong concordance (RMSE = 0.022) between the two alignment technologies, indicating that without modifications, tRNAZAP matches the performance of BWA-MEM (**Figure 7.B**). In addition to maintaining a high level of alignment identity concordance, the tRNAZAP alignment procedure matched the primary tRNA class assignment produced by BWA-MEM for 99.43% of reads (**Figure 7.C**). We repeated this analysis using minimap2 as the non-tRNAZAP aligner. This produced nearly identical results which is not surprising due to the high sequence identity expected from IVT DRS (**Supplementary Figure S2**).

**Figure 7.**
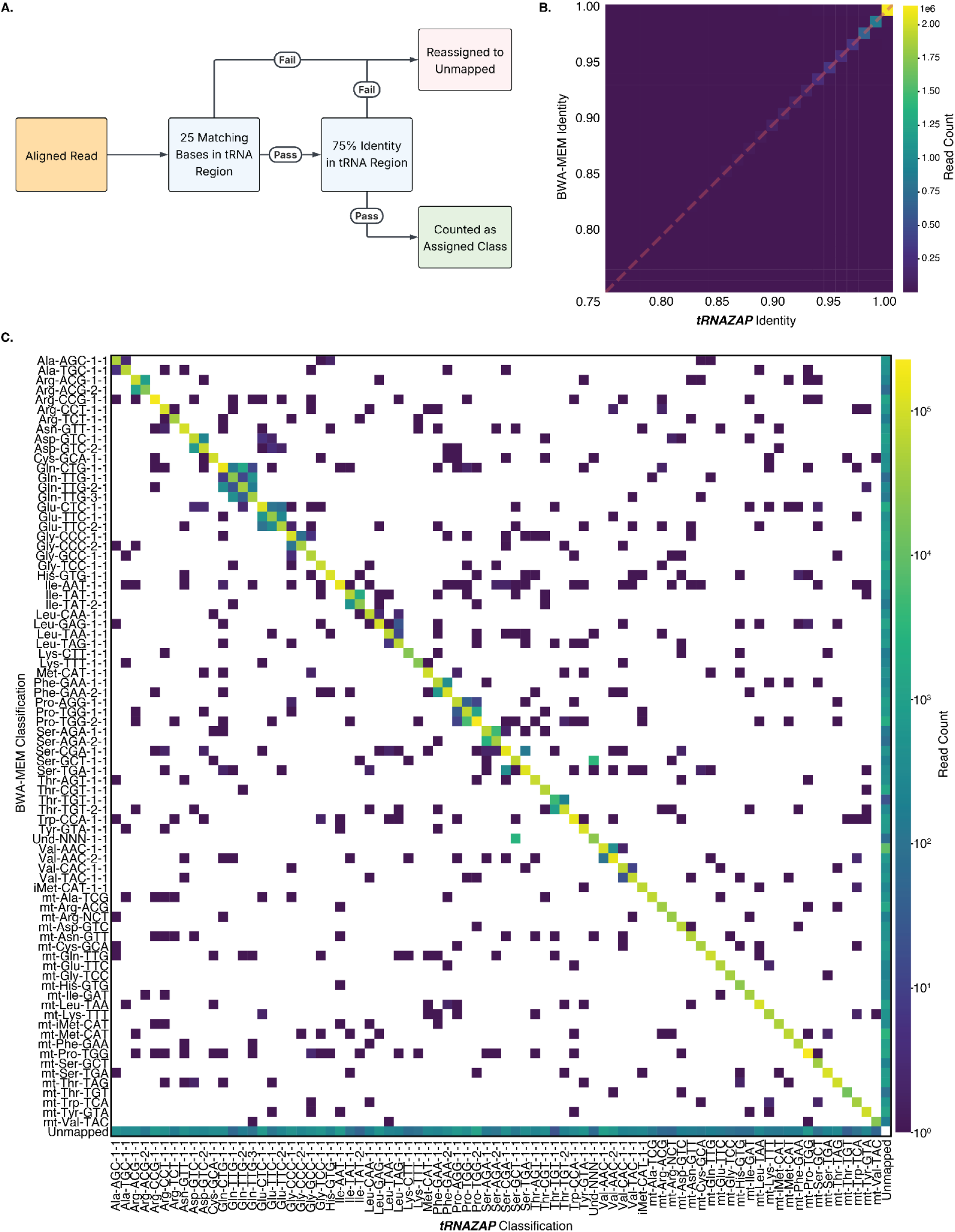
*tRNAZAP* alignment results for IVT *S. cerevisiae* tRNA. **A.** A schematic showing the process by which tRNAZAP removes spurious alignments from model classification. **B**. Comparison of identity between BWA-MEM, the current state-of-the-art alignment method, and tRNAZAP’s alignment method for the *S. cerevisiae* IVT test dataset. **C**. The confusion matrix of classes when BWA-MEM is compared to tRNAZAP, including alignment-based filtering.

### Oligonucleotide pools *tRNAZAP* alignment performance

To further evaluate the performance of tRNAZAP, we generated three mixed oligonucleotide pools containing *S. cerevisiae* and *Escherichia coli* (*E. coli*) tRNA. These pools collectively represent 34 of the 79 annotated *S. cerevisiae* tRNA classes; the remaining 45 classes are absent by design and therefore constituted “unacceptable” targets. An ideal aligner should assign no reads to these absent classes. We used this controlled dataset to compare tRNAZAP with BWA-MEM, quantifying (i) the extent to which the two aligners agree, and (ii) the degree to which each aligner incorrectly maps reads to unacceptable (absent) classes. The IVT tRNA classes present in each pool are listed in **Supplementary Table S4**.

Direct RNA sequencing of these pools produced a total of 61,117,500 reads. All reads were independently processed using both BWA-MEM and tRNAZAP. Because the pools contained transcripts from both species, BWA-MEM was expected to align reads originating from both *S. cerevisiae* and *E. coli*. In contrast, tRNAZAP is designed to identify only *S. cerevisiae* tRNA. Accordingly, we explicitly assessed whether tRNAZAP correctly refrained from assigning *E. coli* tRNA reads, while successfully detecting *S. cerevisiae*-derived tRNA reads.

BWA-MEM successfully aligned 43,730,439 reads. Among these, 30,379,703 reads were assigned to *S. cerevisiae* and 13,350,736 reads were assigned to *E. coli* (**Figure 8.A**). Among the 30,379,703 reads that BWA-MEM aligned to *S. cerevisiae*, tRNAZAP also produced a *S. cerevisiae* alignment for 24,469,939 reads. Within this overlapping set, 97.63% were assigned to the same *S. cerevisiae* tRNA type by both BWA-MEM and tRNAZAP, indicating a high level of concordance at the tRNA-isoacceptor level, while the remaining reads were assigned to a different *S. cerevisiae* tRNA type by tRNAZAP. The remaining 5,909,764 reads in BWA-MEM aligned *S. cerevisiae* read set were not assigned a tRNA by tRNAZAP and were classified as unmapped. **Figure 8.A** presents a summary of the read-level alignment outcomes between BWA-MEM and tRNAZAP, stratified by BWA-MEM and tRNAZAP alignment status. **Figure 8.B** shows the assignment of reads identified as *S. cerevisiae* by at least one of the two aligners to tRNA types, regardless of their presence or absence in the pools.

**Figure 8.**
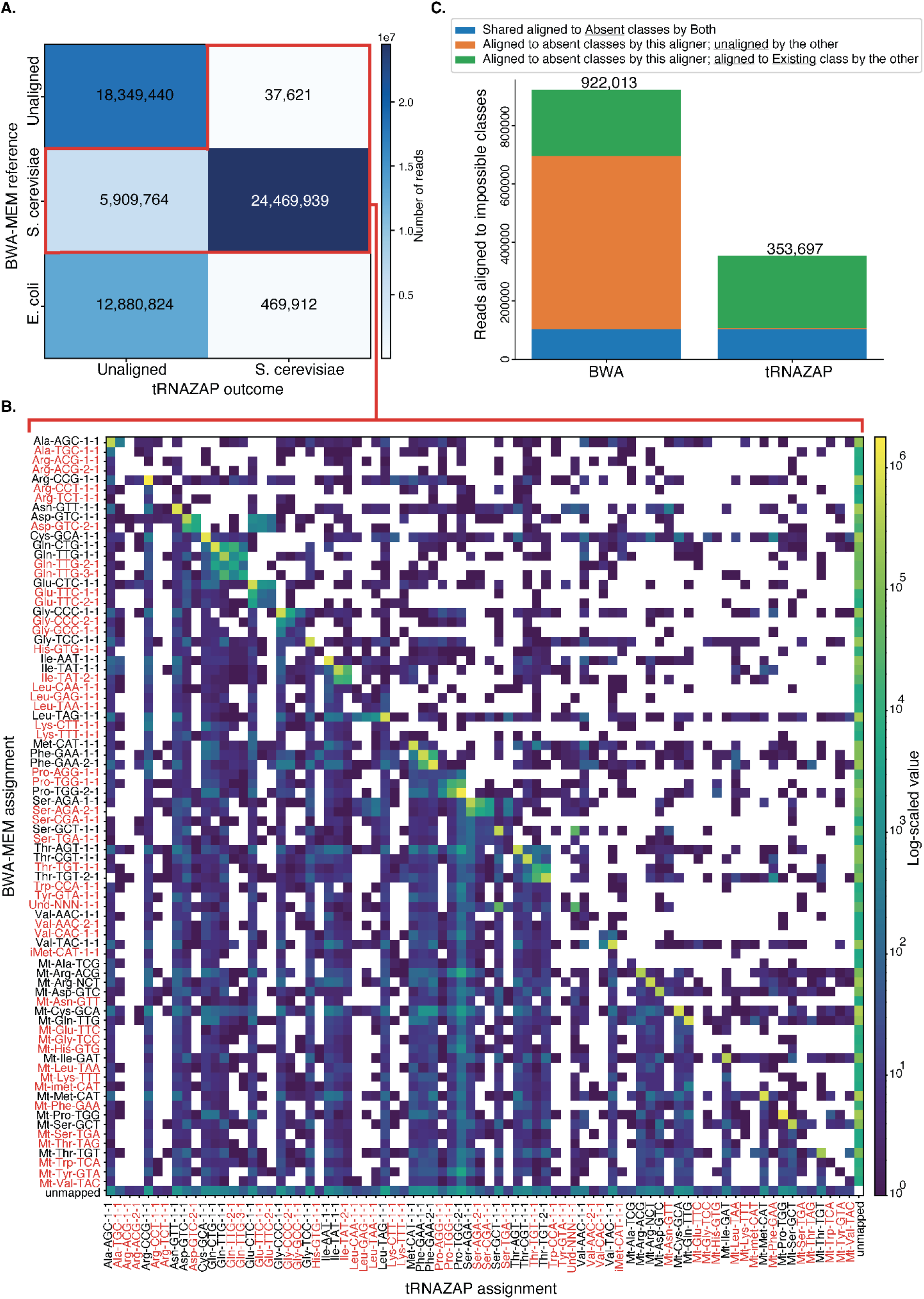
Comparison of alignment outcomes between BWA-MEM and tRNAZAP on mixed *Saccharomyces cerevisiae* and *Escherichia coli* oligonucleotide pools. **A.** Rows indicate the alignments assigned by BWA-MEM (unmapped, *S. cerevisiae*, or *E. coli)*, while columns denote whether tRNAZAP produced an alignment or reported the read as unmapped. Cells are annotated with read counts, illustrating agreement and disagreement between the two methods. **B**. Confusion matrix comparing tRNA type assignments for reads concordantly aligned to *S. cerevisiae* by BWA-MEM and tRNAZAP. Rows represent tRNA types assigned by BWA-MEM, columns represent tRNA types assigned by tRNAZAP, and cell values indicate row-normalized proportions of reads. The red-labeled tRNA classes belong to the Absent classes. **C**. Distribution of read assignments to absent *S. cerevisiae* tRNA classes. Bars summarize how many reads each aligner mapped to classes not represented in the oligonucleotide pools, partitioned into (i) assignments shared by both BWA-MEM and tRNAZAP, (ii) assignments made by one aligner when the other reported the read as unmapped (unaligned), and (iii) assignments made by one aligner when the other mapped the read to an existing class.

Of the 13,350,736 reads aligned to *E. coli* by BWA-MEM, tRNAZAP aligned 469,912 reads to one of the *S. cerevisiae* types, while the remaining 12,880,824 reads were unmapped by tRNAZAP, consistent with its intended species specificity. In total, 18,387,061 reads were unmapped by BWA-MEM. A subset of 37,621 reads was aligned by tRNAZAP, indicating instances where tRNAZAP recovered alignments not detected by BWA-MEM.

We next analyzed alignments to classes that are absent from the oligonucleotide pools and therefore represent unacceptable targets (**Figure 8.C**). BWA-MEM assigned 922,013 reads to these classes, whereas tRNAZAP assigned 353,697, indicating a substantially lower alignment to unacceptable (absent) classes for tRNAZAP. Of these, 103,557 reads were misassigned to absent classes by both aligners. After accounting for these overlaps, BWA-MEM contributed 818,456 unique assignments to unacceptable (absent) classes and tRNAZAP contributed 250,140, demonstrating that the majority of incorrect alignments are aligner-specific rather than shared errors. Directional disagreement between the aligners further highlights their differing behaviors: among the reads BWA-MEM mapped to unacceptable classes, tRNAZAP classified 591,872 as unaligned, whereas only 3,323 reads that tRNAZAP mapped to unacceptable classes were not aligned by BWA-MEM. Notably, 226,584 of BWA-MEM’s unacceptable alignments corresponded to reads for which tRNAZAP identified an acceptable (existing) class, while 246,817 of tRNAZAP’s unacceptable alignments were mapped by BWA-MEM to acceptable classes. **Figure 8.C** shows the distribution of reads mapped to absent classes from the oligonucleotide pools for each aligner, partitioned based on the behavior of the other aligner.

### Runtime Benchmarking of tRNAZAP

To assess the runtime performance of tRNAZAP, we evaluated both the inference and alignment stages using matched subsets of POD5 files and unaligned BAM files containing one million reads. tRNAZAP operates directly on raw nanopore signal data stored in POD5 and unaligned BAM formats, whereas BWA-MEM requires fastq sequences prior to alignment. Fastq files are generated from unaligned BAM files, introducing an extra preprocessing step for BWA-MEM alignment. Using a single NVIDIA V100 GPU (16 GB VRAM) with a batch size of 1,024, tRNAZAP’s inference stage processed the signals in 13 min 25.032 s, and the subsequent alignment stage completed in 2 min 2.138 s, yielding a total runtime of approximately 15 min 27 s. For BWA-MEM, the fastq generation step alone required 1.348s, followed by 4 min 54.354s for alignment.

## Discussion and Conclusion

We present tRNAZAP, a computational toolkit for analyzing tRNA molecules from Nanopore direct RNA sequencing. tRNAZAP consists of three integrated components: a transformer-based sequence-to-sequence model that classifies tRNA type and segments ionic current signals to isolate tRNA regions, a sequence extraction module that uses segmentation boundaries and nanopore Dorado move-tables to obtain query sequences, and an alignment algorithm that maps query sequences to their predicted references using an augmented Needleman-Wunsch approach. tRNAZAP leverages direct interpretation of ionic current signal for classification of tRNA. This circumvents the pitfalls of sequence-based classification, which can be complicated by an intermediate basecalling model that is presently not trained for tRNA.

tRNAZAP models were trained using *in vitro* transcribed tRNA data. We documented ∼98% classification accuracy on yeast tRNA classes and segmentation of signal boundaries with an IoU of ∼0.98. The pairwise alignment strategy achieved a median alignment identity of 98.67%.

As the current models are trained on IVT data, fine-tuning on biological samples with native modifications will be necessary before drawing any biological conclusions^25^. The ionic current signal analysis strategy we implemented for tRNAZAP is generalizable beyond tRNA. It could be extended to any class of DNA or RNA (e.g., other small RNA species) that can be adapted on the 5’ and 3’ ends. An attractive possible application of this methodology could be for microRNA, a class of short RNAs with extensive documentation on their regulatory role.

Ionic current signal readouts include sequence and modification information from nucleic acid strands as they are translocated through a nanopore. There is a gap in tools for addressing analysis challenges associated with short, heavily modified molecules like tRNA. tRNAZAP is a generalizable toolkit for segmentation and classification of Nanopore ionic current signals, and for more precise Nanopore tRNA sequence alignments.

## Key points

- ***tRNAZAP*** segments tRNA regions from Nanopore ionic current with ∼0.98 IoU.
- ***tRNAZAP*** classifies Nanopore tRNA reads for 79 yeast tRNA classes with ∼98% accuracy.
- ***tRNAZAP*** alignment achieves 98.67% median identity for IVT tRNA.
- ***tRNAZAP*** uses Nanopore ionic current signals and achieves tRNA classification independent of basecalling accuracy.

## Data and Code Availability

All the codebase, models, and configuration used in this study are publicly available in the GitHub repository (https://github.com/genometechlab/trnazap) with instructions on how to use them. Sequence-level data will be publicly available through the European Nucleotide Archive.

