## Supplementary Information for "*tRNAZAP*: Deep learning toolkit for nanopore tRNA signal analysis"

### Supplementary Figures

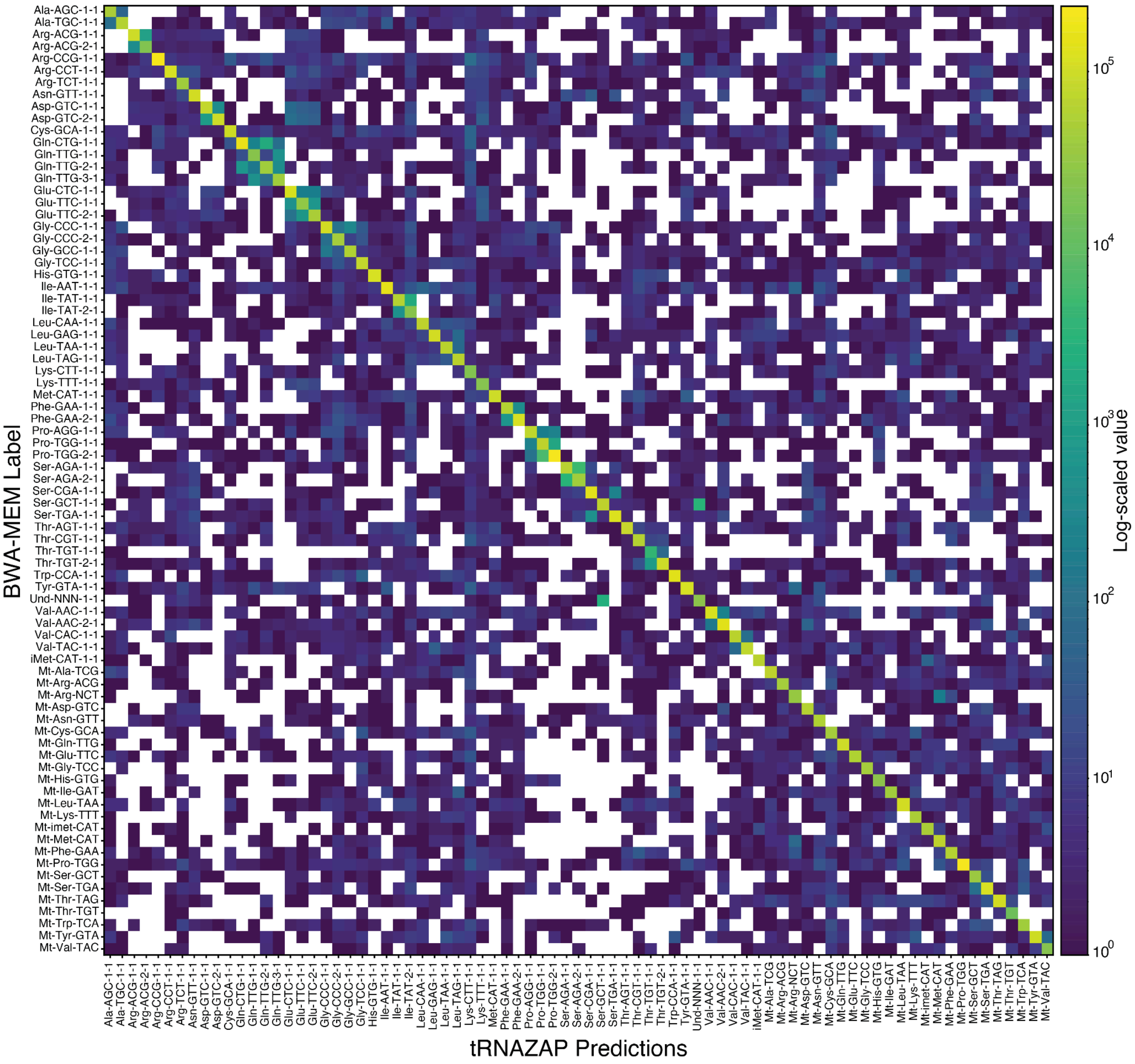


**Supplementary Figure S1.** Confusion matrix showing classification across 79 yeast tRNA classes. Each cell reports the number of reads assigned to the BWA-MEM class on the y-axis and the tRNAZAP class on the x-axis. Counts are displayed on a logarithmic color scale (LogNorm) to accommodate the wide dynamic range, with zero-count cells masked in white.

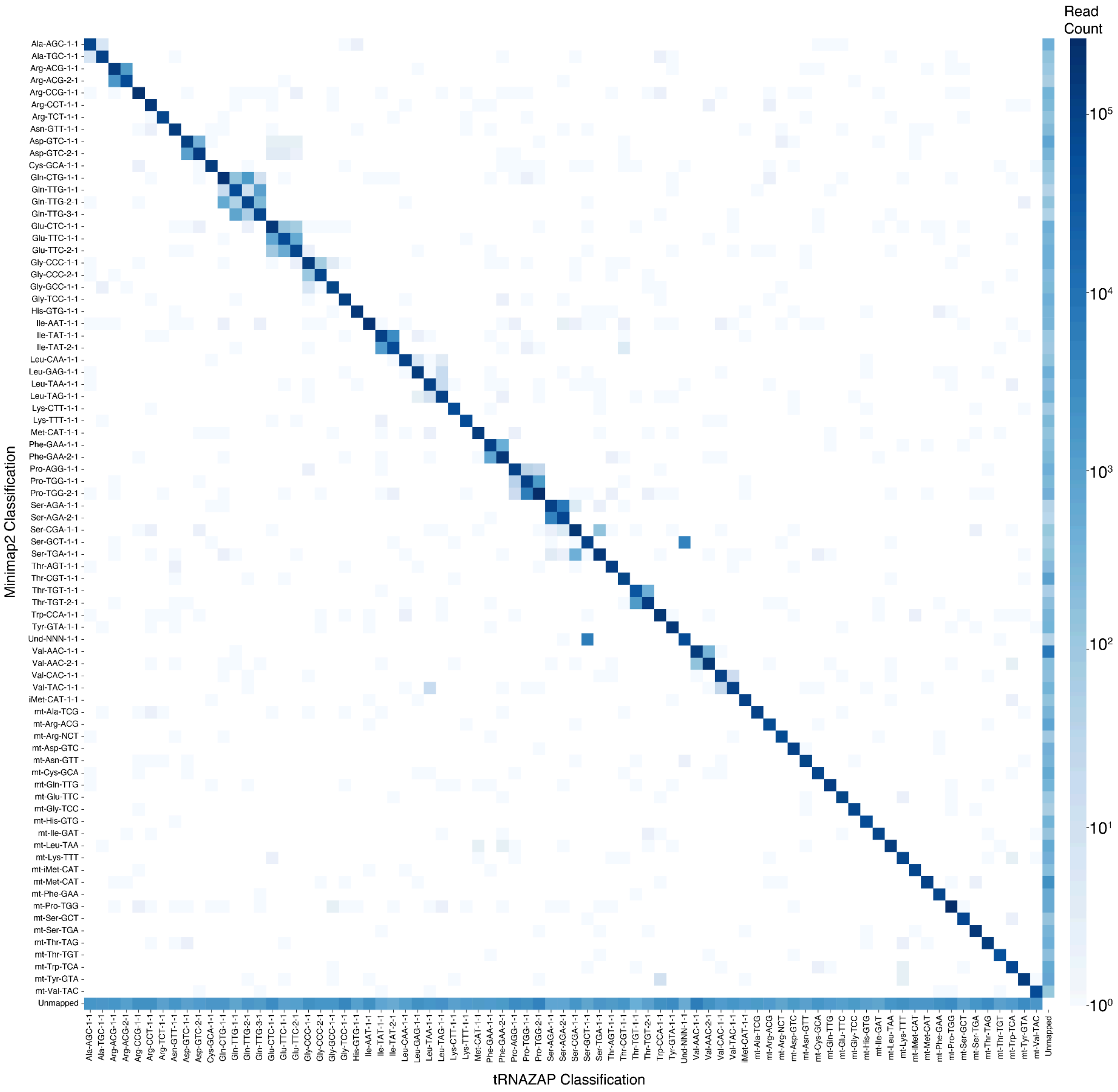


**Supplementary Figure S2.** Confusion matrix with the tRNAZAP vs. minimap2 (-x map-ont) alignment classifications. tRNAZAP prediction classes are depicted on the x-axis while minimap2 prediction classes are decpicted on the y-axis. The color of each cell represents the number of aligned reads for that combination of tRNAZAP and minimap2 classification.
